# Multi-omics analysis reveals the regulatory mechanism of flavonol biosynthesis during the formation of petal color in *Camellia nitidissima*

**DOI:** 10.1101/2023.01.03.522545

**Authors:** Yi Feng, Jiyuan Li, Xian Chu, Hengfu Yin, Zhengqi Fan, Suhang Yu, Minyan Wang, Xinlei Li, Weixin Liu

**Affiliations:** Key Laboratory of Tree Breeding of Zhejiang Province, Research Institute of Subtropical Forestry, Chinese Academy of Forestry, Hangzhou, Zhejiang, China; Jinhua Moxian Horticultural Engineer Co. Ltd., Jinhua, Zhejiang, China

**Keywords:** *Camellia nitidissima*, Petal color, Flavonol, Metabolome, Proteome, Transcriptome, Hormone

## Abstract

*Camellia nitidissima* is a prized woody flower with golden-yellow flowers, and has high ornamental, medicinal and economic value. Previous works indicated that the content of flavonol accumulated greatly during golden petal formation. However, the molecular mechanism of golden flower formation in *C. nitidissima* remains largely unknown. In this study, we performed an integrative anlysis of transcriptome, proteome and metablome of petals at five developmental stages to construct the regulatory network during golden flower formation in *C. nitidissima*. Metablome anlysis showed that two flavonols, the quercetin and kaempferol glycosides, were highly accumulated in the golden petals. Furthermore, transcriptome and proteome sequencing suggested that the expression of flavonol biosynthesis genes or proteins was increased in golden petal stage, whereas expression of anthocyanin and proanthocyanidin genes or proteins were decreased. Six *MYB* and 20 *bHLH* genes were identified as potentially involved in flavonoid biosynthesis, and the brassinosteroid (BR) and jasmonate (JA) pathways were positively and negatively corretated with flavonol biosynthesis, respectively. Network correlation analysis suggested close relationships among BR and JA, MYB and bHLH, and the flavonoid pathway and flavonoid metabolites. Overall, this study shows a potential regulatory mechanism of flavonol biosynthesis duing golden petal formation in *C. nitidissima*.

**Highlight:** The BR and JA pathway may positively and negatively regulate flavonol synthesis in *Camellia nitidissima*, respectively.

## Introduction

Flavonoid is the largest pigment group in plants and the key factor of petal color formation in most plants (Noda, 2018; Zhao *et al*., 2022). According to their structure, flavonoids can be classified into flavonols, anthocyanins, proanthocyanidins, flavones, isoflavones, chalcones, aurones and so on (Liu *et al*., 2021). Anthocyanins can give plants red, orange, purple and blue (Zhang *et al*., 2014); and flavonols, chalcones and aurones are the important pale-yellow or yellow pigments in plants (Hoshino *et al*., 2019; Yang *et al*., 2020). Besides, flavonoids can be as edible pigments and taste-regulating components in food and wine (Winkel-Shirley, 2001; Yang *et al*., 2014), and are also associated with health-promoting functions such as antioxidant, vasodilator, anti-carcinogenic, anti-aging activities (Nabavi *et al*., 2018; Wang, ZL *et al*., 2018).

Flavonoids are synthesized through the phenylpropanoid pathway, and more than 9000 flavonoids have been identified in plants (Sun *et al*., 2020b; Liu *et al*., 2021). Flavonols are an important branch of the flavonoid biosynthesis pathway. The biosynthesis of flavonols begins with phenylalanine, which is converted to coumaroyl-CoA by 4-coumarate: CoA ligase (4CL), cinnamic acid 4-hydroxylase (C4H), and phenylalanine ammonia lyase (PAL) (Winkel-Shirley, 2001; Wohl & Petersen, 2020). Dihydroflavonols, which include dihydroquercetin (DHQ), dihydrokaempferol (DHK), and dihydromyricetin (DHM), are generated from coumaroyl-CoA under the catalysis of chalcone isomerase (CHI), chalcone synthase (CHS), flavanone 3’-hydroxylase (F3’H), flavanone 3-hydroxylase (F3H), and flavanone 3’5’-hydroxylase (F3’5’H) (Nakatsuka *et al*., 2014; Si *et al*., 2022). Dihydroflavonols are the key intermediate metabolites in flavonoid biosynthesis and can be converted to flavonols through the bioactivity of flavonol synthase (FLS) (Jiang *et al*., 2020). In addition, anthocyanin and proanthocyanidin can be generated from dihydroflavonols by dihydroflavonol 4-reductase (DFR), anthocyanidin synthase (ANS), leucoanthocyanidin reductase (LAR), anthocyanidin reductase (ANR) (Giampieri *et al*., 2018). DFR and FLS compete with the substrate dihydroflavonol and flowe into the anthocyanin, proanthocyanidin and flavonol biosynthetic pathways, respectively (Luo *et al*., 2016).

In the transcriptional regulation of flavonol metabolism, MYB transcription factors and their MBW protein complex, composed of bHLH, MYB, and WD40, are the most widely and clearly studied factors (Xu *et al*., 2015; Zhang *et al*., 2021). In *Arabidopsis thaliana*, subgroup 7 MYB family members AtMYB111, AtMYB12, and AtMYB11 activate the expression of *FLS, F3H, CHS*, and *CHI* genes, leading to increased flavonol content (Stracke *et al*., 2007). CsMYB60, a homologous protein of AtMYB111, induces the expression of *CsLAR* and *CsFLS* through binding to their promoters and increases the expression of *Cs4CL* and *CsCHS*, thereby promoting the accumulation of proanthocyanidins and flavonols in *Cucumis sativus* (Li *et al*., 2020). In addition to MYB transcription factors, hormones including jasmonate (JA), auxin, ethylene, and gibberellin (GA) are involved in the regulation of flavonol metabolism in plants. Auxin and ethylene differentially increase the expression of flavonol biosynthesis genes and the biosynthesis of flavonol (Lewis *et al*., 2011). In *A. thaliana*, GA inhibited flavonol accumulation through DELLA proteins, the core negative regulator of the GA signaling pathway; and RGA and GAI, two DELLA proteins, interacted with MYB12 and MYB111 to regulate the expression of flavonol biosynthesis genes and promote flavonol accumulation (Tan *et al*., 2019). The preharvest methyl jasmonate (MeJA) treatments has been shown to increase the content of flavonlos such as myricetin and quercetin and the acitivity of PAL enzyme (Flores & Ruiz del Castillo, 2014).

The number of Camellia species and varieties has exceeded 20000, but its petal colors are mainly red, and yellow flower varieties are rare, accounting for less than 1% (Guan *et al*., 2014). *Camellia nitidissima* is prized woody flower, known for its unique golden-yellow petal color in Camellia (He *et al*., 2017), and has high ornamental and economic value. Besides, the golden-yellow petals of *C. nitidissima* are rich in flavonoids, especially flavonols (Tanikawa *et al*., 2008; Zhou *et al*., 2017; Liu *et al*., 2022), which have a variety of physiological activities including antioxidant, anti-aging, lipid lowering, blood pressure lowering effects and have huge economic value in medical care and food production (Nabavi *et al*., 2018; Wang, ZL *et al*., 2018). Therefore, *C. nitidissima* is a precious species integrating ornamental, medicinal and edible functions, as well as a valuable resource for molecular mechanism research of yellow petal formation and yellow camellia breeding.

Previous works suggested that flavonols were the main pigment in the golden-yellow petals of *C. nitidissima* (Tanikawa *et al*., 2008; Liu *et al*., 2022). However, the molecular mechanism of flavonol biosynthesis during the formation of golden color in *C. nitidissima* petals remains largely unknown. To investigate this, we used petals of *C. nitidssima* at five different development periods as study materials and carried out combined multi-omics analyses (metabolome, transcriptome, and proteome) to identify the potential pathways of flavonol accumulation during gloden flower formation in *C. nitidssima*.

## Materials and methods

### Plant materials

The *C. nitidissima* plants were from the Germplasm Resources of Camellia (Qianjia village, Fuyang city, Zhejiang province, China), where they have been grown in the field, and were about 15 years old. Petals were harvested at five different developmental stages, early-bud stage (S0), mid-bud stage (S1), late-bud stage (S2), half-opening stage (S3), and complete-opening stage (S4), harvested in February 2021.

### Metabolic analysis

The metabolites were qualitatively quantified by ultra-performance liquid chromatography (UPLC)–electrospray ionization (ESI)–tandem mass spectrometry (MS/MS). Qualitative analysis of metabolites was performed based on the Metware Biotechnology Self-built Database (Wuhan, China) and secondary spectral information. Multiple reaction monitoring analysis with triple four-stage MS was used for metabolite quantification. Unsupervised PCA and OPLS-DA (partial least squares discriminant analysis) were performed to analyze metabolites. The analysis of hierarchical clustering in different samples was performed using R software (www.r-project.org/). To identify differential expressed metabolites (DEMs), the metabolome data were analyzed using the OPLS-DA model. The criteria, a fold change ≥2 or fold change ≤0.5 based on the OPLS-DA results and variable importance in project ≥1, was used for significantly differential metabolites identification. And the analysis of KEGG (Kyoto Encyclopedia of Genes and Genomes) of significantly differential metabolites was performed, subsequently.

### RNA sequencing (RNA-seq) analysis

Total RNA was extracted from the flowers of *C. nitidissima* using an RNA Prep Pure kit for plants (Tiangen, Beijing, China). The library of RNA-seq was generated and sequenced on an Illumina HiSeq platform (Illumina, San Diego, CA, USA). Gene expression levels were estimated by RSEM (Li & Dewey, 2011), and the expression abundance of the corresponding unigenes was calculated by the Fragments per Kilobase of Transcriptome per Million Mapped Reads method. Gene function was annotated based on the following databases: KEGG; KOG/COG (COG: Clusters of Orthologous Groups of Proteins; KOG: euKaryotic Ortholog Groups); GO (Gene Ontology); Nr (NCBI non-redundant protein sequences); Trembl (a variety of new documentation files and the creation of TrEMBL). Differential expression analysis between two groups was executed through DESeq2 software (Leng *et al*., 2013). Benjamini and Hochberg’s method were carried out for the correction of P-values, and the posterior probability values were used for the correction of false discovery rate (FDR). The thresholds of significant differential expression were FDR < 0.05 and |log_2_ (foldchange)| ≥ 1.

### Protein sample preparation and MS detection and data analysis

Protein in solution was precipitated using acetone, and the ground flower tissue was mixed with four volumes of lysis buffer (8 M urea, 100 mM Tris-Cl, 10 mM dithiothreitol) and incubated at 37 °C for 1 h. Subsequently, 40 mM iodoacetamide was added to the complex solution. The Bradford method was used to determine protein concentration. After protein quantification, 50-μg samples were separated by sodium dodecyl sulfate polyacrylamide gel electrophoresis, and the protein bands were stained with Kemas Brilliant Blue. The extracted proteins were reduced and alkylated and then digested with trypsin. Peptides were desalted using a Sep-Pak C18 column and vacuum dried.

The MS data were acquired using a Q Exactive HF-X mass spectrometer in tandem with an EASY-nLC1200 liquid-phase liquid chromatography system. DIA-NN software was used to establish a spectrum library based on the protein sequence database of *Camellia sinensis* var. *sinensis* in the UniProt database; protein identification was performed, and quantitative information was extracted. The test results were screened using a threshold of 1% FDR. The quantification intensity information obtained from DIA analysis was used for difference comparison, with t-test analysis after log2 transformation, data filling (imputation algorithm in Perseus software), and data normalization: fold change ≥1.5, P < 0.05 indicated upregulated proteins, fold change ≤1/1.5, P < 0.05 indicated downregulated proteins.

### Interaction network analysis

Pearson’s correlation coefficients were used for the establishment of interaction network, and R environment (https://www.r-project.org/) was performed for the calculation of Pearson’s correlation coefficients; correlations with a coefficient of R ≥ 0.8 or R ≤ –0.8 and P ≤ 0.05 were retained. The relationships between candidate genes, including transcription factors (TF) genes and structural genes, proteins, and flavonoid components were visualized using Cytoscape (v. 3.9.2).

### Quantitative real-time PCR

To validate the RNA-seq results, 14 flavonol- and hormone-related genes were selected for quantitative real-time PCR (qRT-PCR) assay. All primers of qRT-PCR were designed using NCBI online, and *GAPDH* was used as the internal normalization gene (Supplementary Table 1). The analysis of qRT-PCR was performed as described previously (Liu *et al*., 2022).

## Results

### Changes in the phenotypic characteristics of developing *C. nitidissima*

The phenotypic changes of petals during five developmental stages were observed (Fig.1). In S0, buds are tiny and tightly closed, and petals are white and nearly translucent. S1 is the mid-bud period, where the bud is more inflated than in S0, and the petals appear white. The S2 is the late-bud stage, when the bud reaches its maximum size and the petals take on a yellow-green color. The flower appears half-opened at S3, revealing the complete floral organs (pistils, stamens, sepals, and corolla), and the corolla has a golden yellow color. The petals are fully expanded at S4, with little change in petal color compared with S3. During the formation of golden color of petals in *C. nitidissima*, flavonoids, especially flavonols, are significantly accumulated (Liu *et al*., 2022). Based on the phenotypic differences, the five developmental stages of petals were further subjected to metabolome, transcriptome, and proteome analysis.

**Fig. 1.**
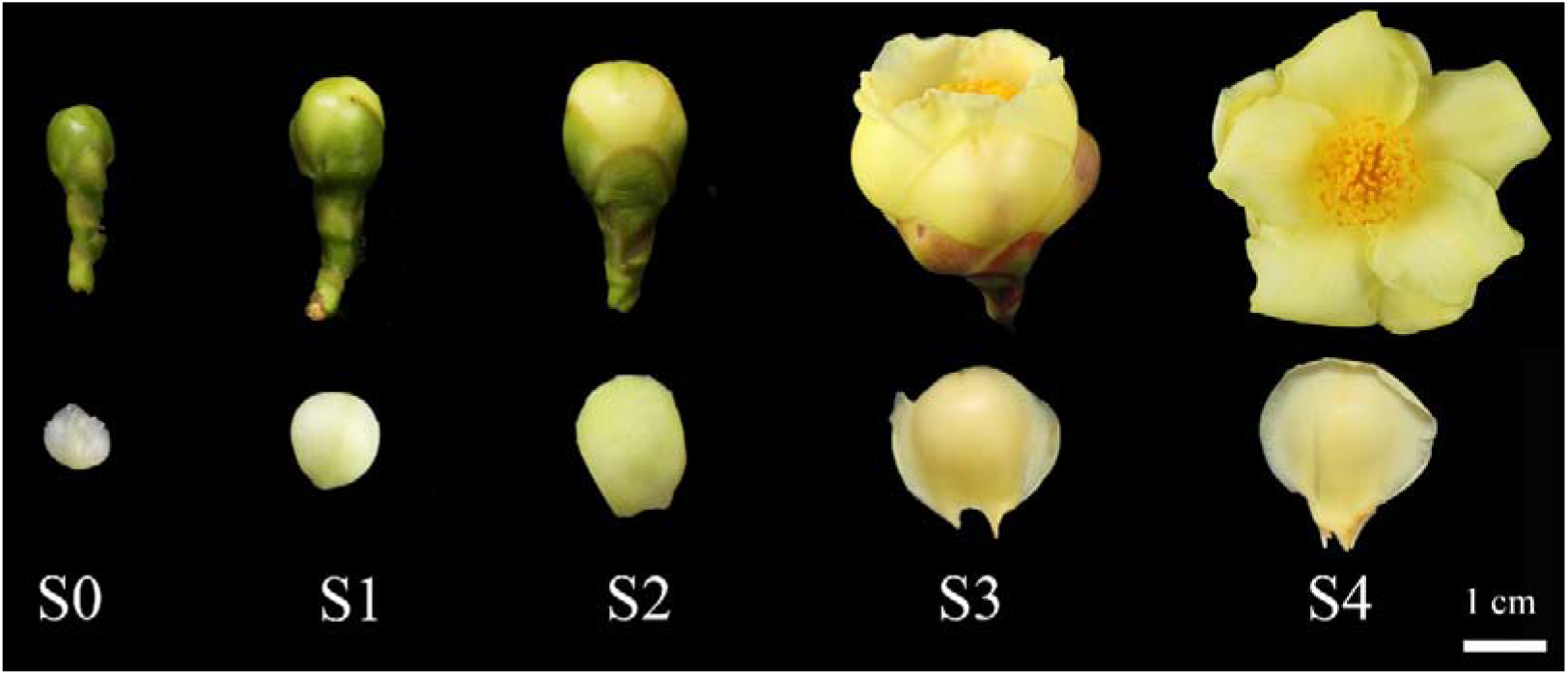
Five developmental stages of petals in *C. nitidissima*. S0, early-bud stage; S1, mid-bud stage; S2, late-bud stage; S3, half-opening stage; S4, complete-opening stage.

### Overview of the metabolome Data

To analyze the metabolic changes of flavonoids in petals of *C. nitidissima*, we performed metabolome annlysis with UPLC-MS/MS. Fifteen samples were divided into five groups with three replicates each for metabolic study. The OPLS-DA score map showed good variability among groups (Fig. 2a). Finally, a total of 323 flavonoids were screened, including 114 flavonols (35.29 %), 59 tannin (18.27 %), 28 flavones (8.67 %), 24 flavanols (7.43 %), 23 flavonoid carbonside (7.12 %), 21 flavanones (6.50 %), 13 anthocyanins (4.02 %), 11 isoflavones (3.41 %), 10 proanthocyanidins (3.10 %), 10 chalcones (3.10 %), eight flavanonols (2.48 %) (Fig. 2b).

**Fig. 2.**
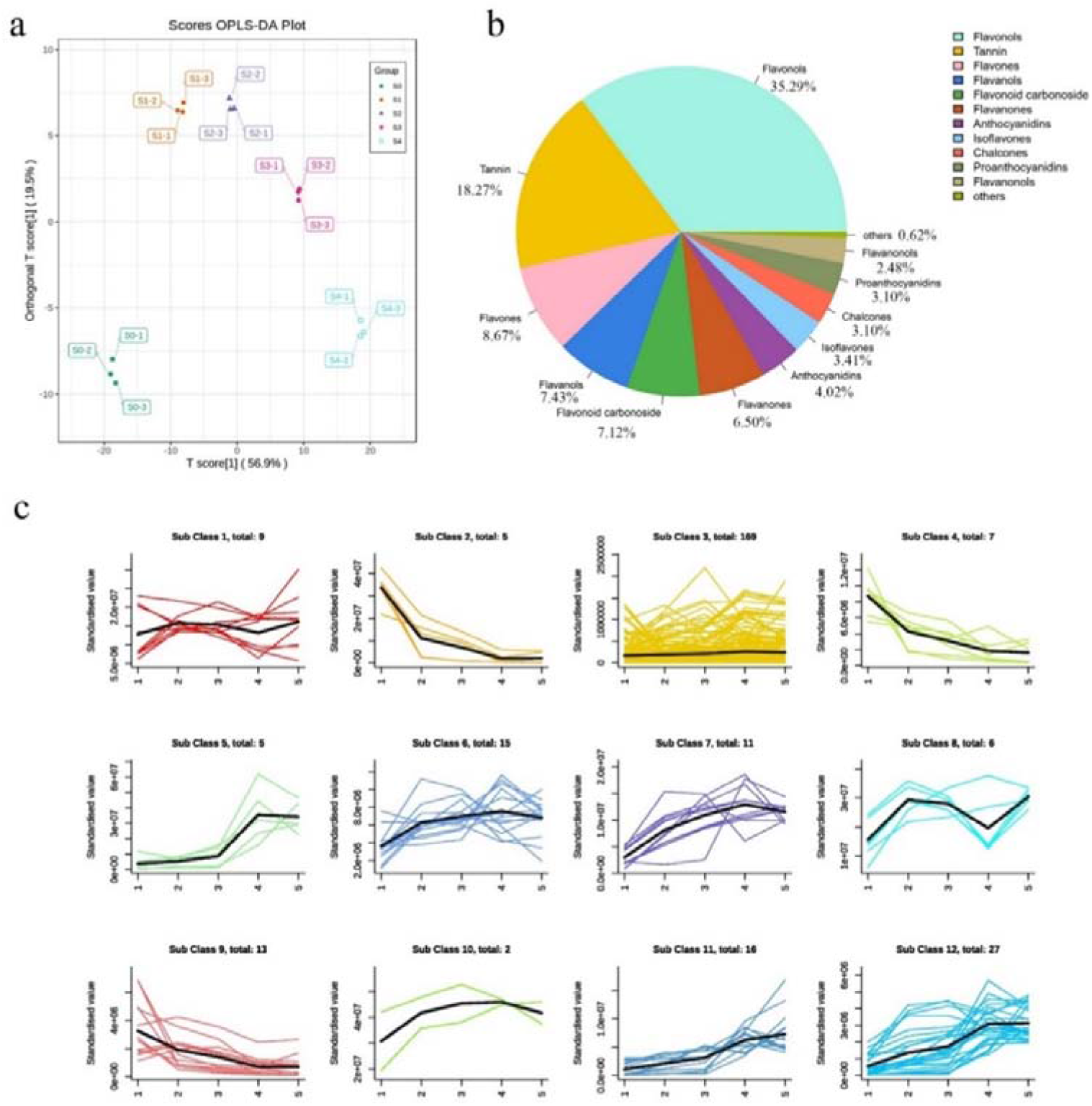
Metabolome analysis of *C. nitidissima*. (a) OPLS-DA Score Map. Each point in the graph indicates a sample, and samples of the same group are represented with the same color. (b) Number of differential metabolites. (c) Expression trends of the identified DEMs in five developmental stages.

### Analysis of the differentially expressed metabolites (DEMs)

We conducted further analysis of the DEMs in the different five developmental stages of petals, a total of 285 DEMs were obtained (Supplementary Fig. 1). K-means clustering analysis was performed to classify the expression patterns of DEMs, 12 sub-classes were obtained and the number ascribed to each class was also recorded (Fig. 2c). Further thermogram analysis showed that there were 97 metabolites positively related to the formation of golden flowers. These included 79 flavonols, in particular, quercetin-related glycosides and kaempferol-related glycosides (Fig. 3a). There were 45 metabolites negatively related to the formation of golden flower, including eight proanthocyanidins and four anthocyanins (Fig. 3b). These results indicate that the accumulation of flavonols glycosides may be key metabolites during the formation of the golden color of *C. nitidissima*. Therefore, to explore the possible mechanism by which flavonols accumulation deiierences in *C. nitidissima*, we conducted transcriptome and proteome sequencing.

**Fig. 3.**
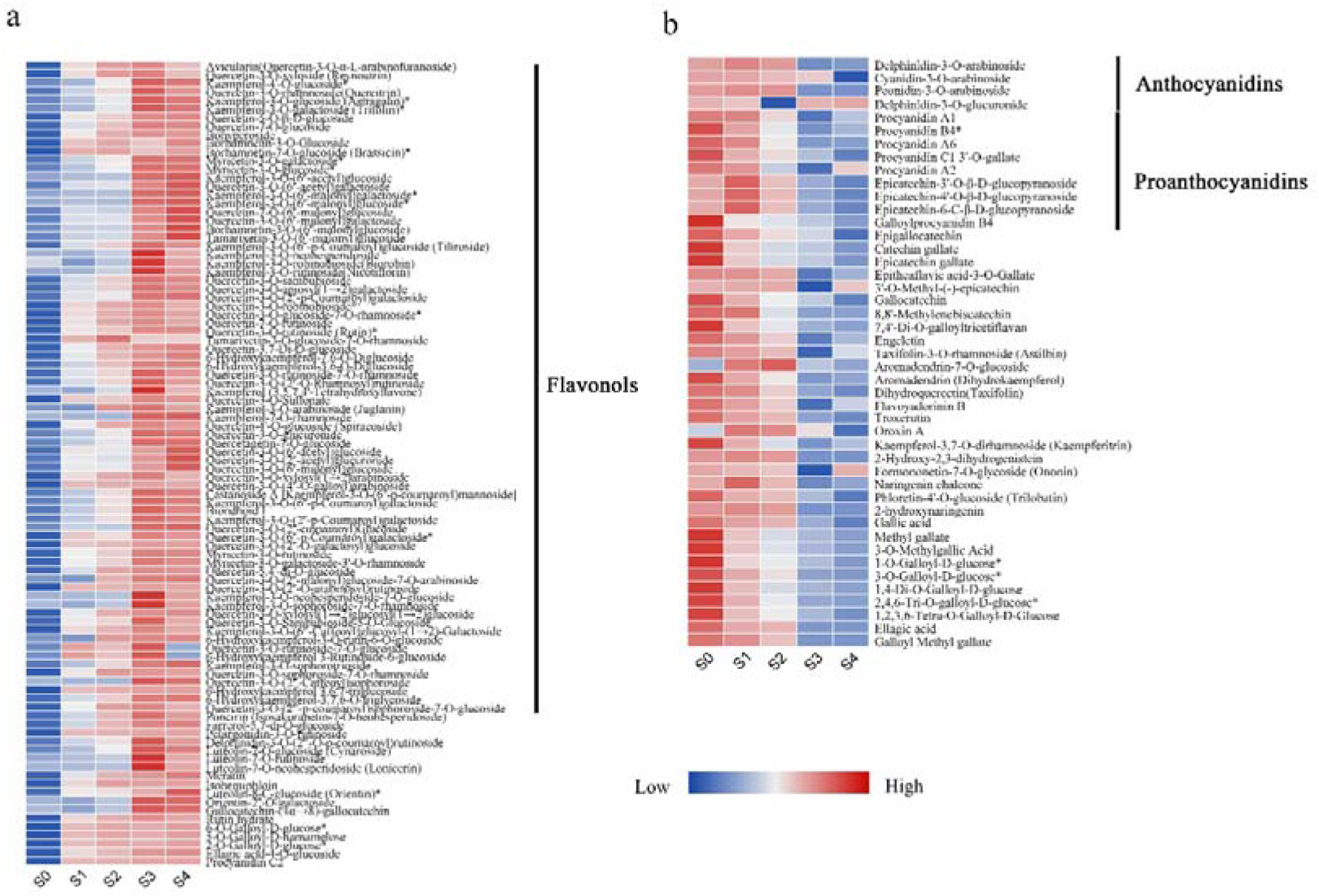
Heat map of DEMs. Heat maps of positively (a) and negatively (b) correlated metabolites during the the formation of golden flowers.

### RNA-seq analysis

RNA-seq was performed on the same 15 samples used for metabolome analysis with an Illumina HiSeq platform. A total of 112.88 Gb clean data were generated after quality filtering. The clean data of 15 samples was 6.37 Gb or more, reached 6.37 −8.32 Gb, with an average of 7.53 Gb (Supplementary Table 2). And the percentage of Q30 base was not less than 92.39 %, and the content of GC reached 44.30-44.69 %. A total of 122,201 unigenes were generated after sequence assembly; the average length of a unigene was 2219 base pairs (bp), with an N50 value of 2699 bp. To annotate the functions of these unigenes, their sequences were submitted to seven functional databases (KEGG, NR, Swiss-Prot, GO, COG/KOG, Trembl, and Pfam) to search for annotations. The number of annotated unigenes in the seven databases ranged from 66,946 to 107,850, corresponding to annotation percentages of 54.78 % to 88.26 % (Supplementary Table 3). Among them, 21,920 (20.14 %) unigenes had good matches with sequences of *Vitis vinifera*, and 4892 unigenes showed high similarity with genes from *Camellia sinensis*, followed by *Vitis vinifera* (Supplementary Fig. 2).

To verify the reliability of the RNA-seq results, 14 flavonoid- and hormone-related genes were carried out for further qRT-PCR investigation. The expression trend of these genes obtained by qRT-PCR was consistent with the results of the RNA-seq (Supplementary Fig. 3), suggesting that RNA-seq data were trustworthy.

### Differentially expressed genes (DEGs) analysis

Correlation analysis of the transcriptome samples was performed for each period, base on the expression levels of the DEGs. The results showed good reproducibility between samples (Figure 4a). DEGs were screened out using the criteria |log2Fold Change| ≥ 1 and FDR < 0.05 thresholds. In total, 46,637 DEGs were identified using the DESeq2/ edgeR package. In the comparisons of S0 vs. S1, S1 vs. S2, S2 vs. S3, and S3 vs. S4, there were 2902, 881, 26,616, and 22,028 DEGs, of which 1234, 423, 12,833, and 12,035 were upregulated and 1668, 458, 13,783, and 9993 were downregulated, respectively (Figure 4b). K-means clustering analysis was carried out to classify the expression patterns of DEGs, and eight sub-classes were identified (Figure 4c). The results suggested that sub-class 6 was relevant to the flavonol accumulation in *C. nitidissima*, whereas the expression of genes in sub-classes 4 and 7 was negatively correlated with flavonol accumulation (Figure 4c).

**Fig. 4.**
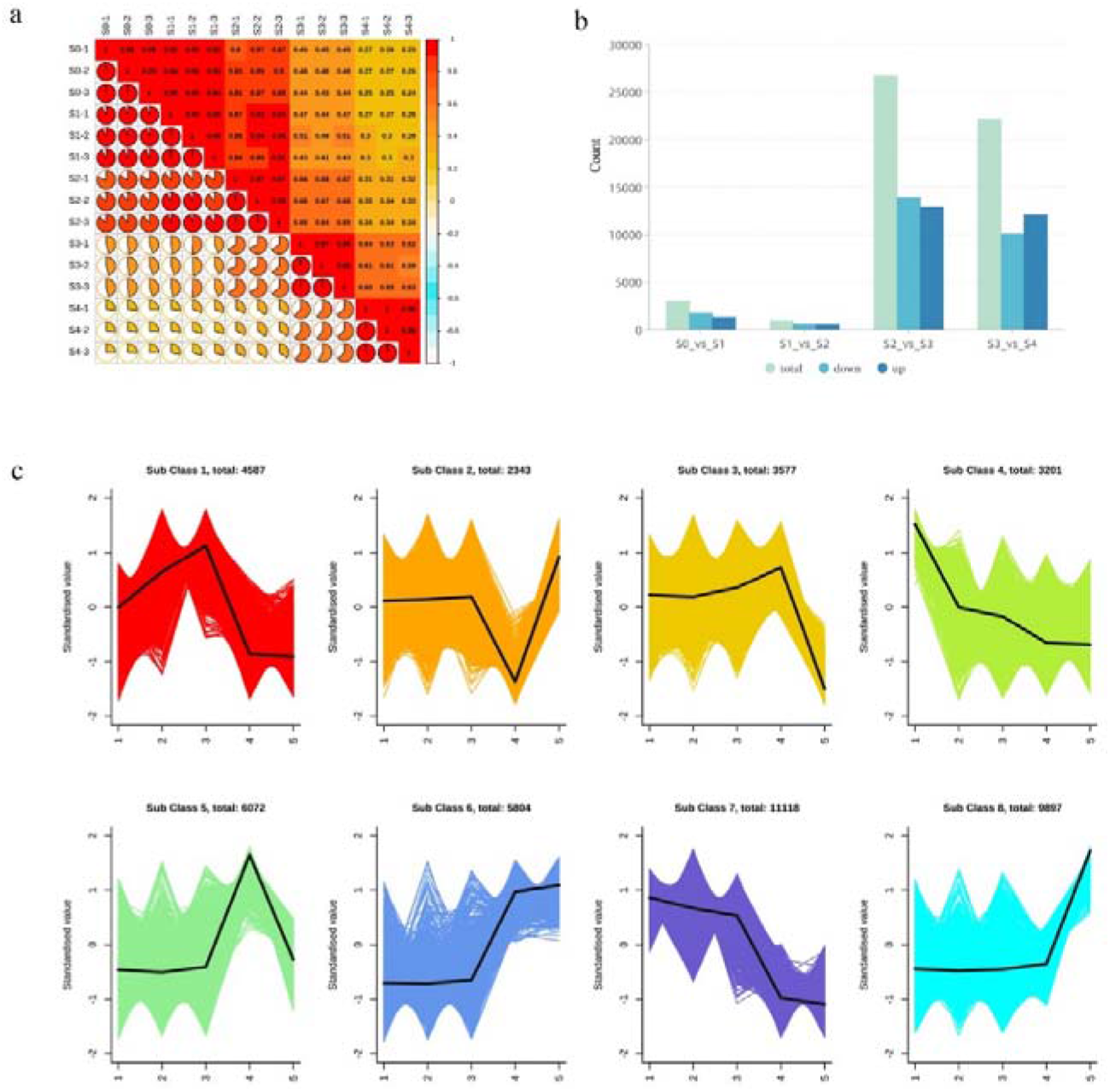
Transcriptome data quality testing. (a) Transcriptome sample correlation analysis. (b) Up-and downregulated unigenes in different comparisons. (c) Expression trends of the identified DEGs in five developmental stages.

We detected many DEGs involved in the flavonoid biosynthesis pathway (Figure 5a), including the biosynthesis genes *PAL*, *C4H, 4CL, CHI, F3H, FLS, DFR*, and *ANS*. Many *MYB* and *bHLH* genes were obtained (Supplementary Fig. 4), of which 20 *bHLH* (Figure 5b) and six *MYB* (Figure 5c) genes were annotated in the flavonoid pathway. Many hormone-related genes, in particular, brassinosteroid (BR) and JA, were screened (Figure 5d, e). These included BR signal transduction genes *BRI1s, BAK1s, BSKs, BSU1s, 14-3-3s, PP2As* and *BZR1s;* JA biosynthesis genes *LOXs, AOSs, OPRs, CYP94B3*, and *CYP94C1s;* and JA signaling genes *JAZs*. These results indicate that flavonoid-related, hormone-related (BR and JA), and *MYB* and *bHLH* genes may have key roles in flavonol biosynthesis in petals of *C. nitidissima*. Network correlation analysis showed that hormone-related (BR and JA), transcription factors (MYB and *bHLH*), flavonoid biosynthesis genes, and flavonoid metabolites were closely related (Fig 6).

**Fig. 5.**
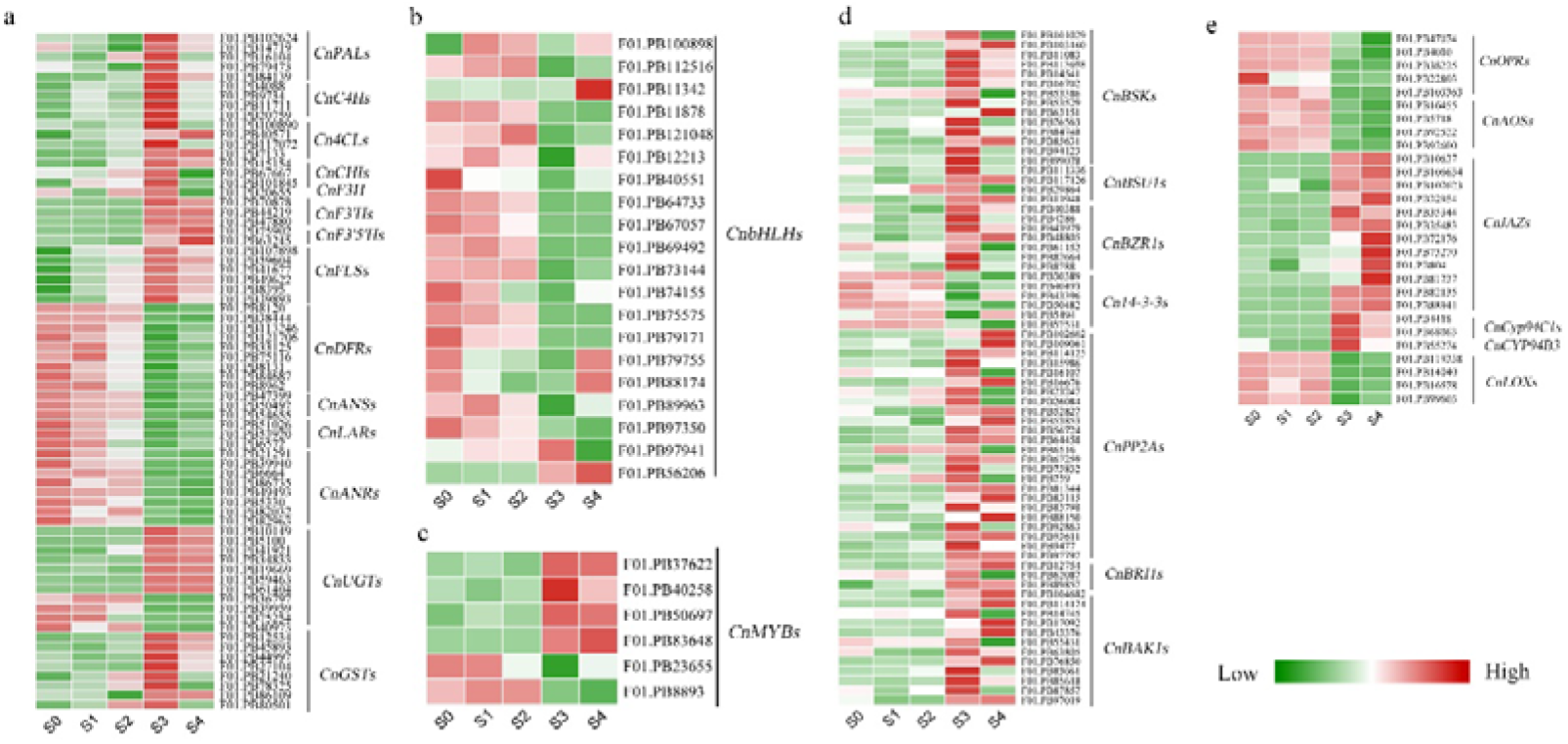
Heat maps of flavonoid (a), bHLH (b), MYB (c), BR-related (d) and JA-related genes (e).

**Fig. 6.**
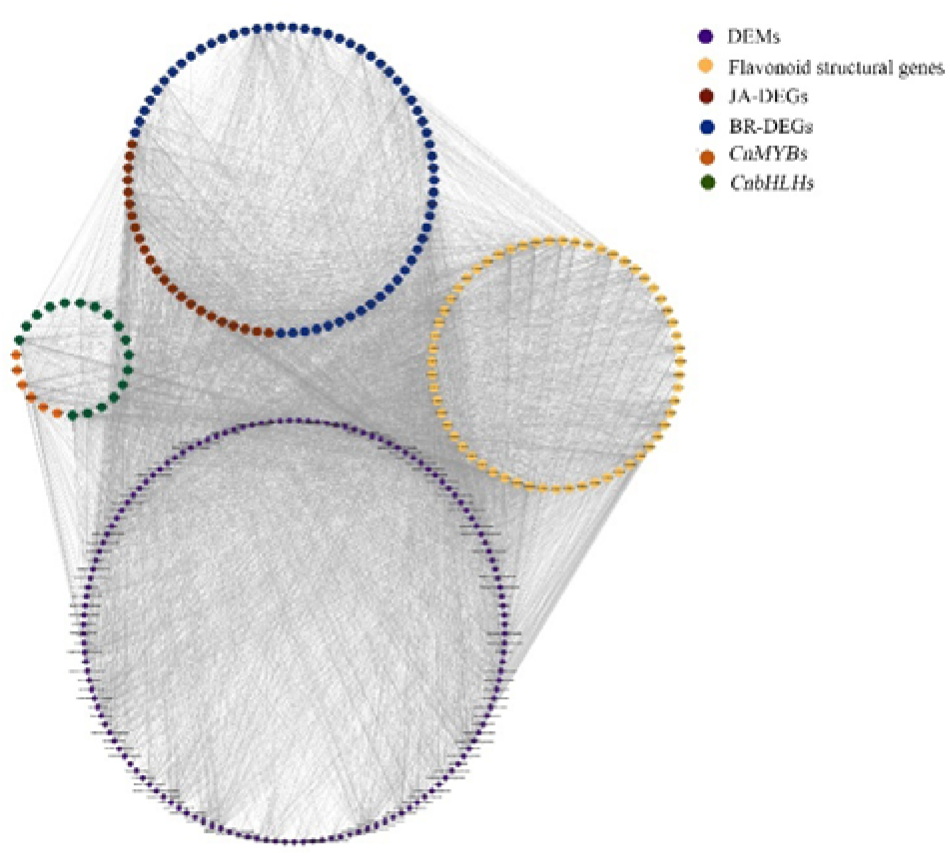
Network correlation analysis of *bHLHs*, *MYBs*, hormone- and flavonoid-related genes, and flavonoid metabolites.

### Proteome sequencing analysis

Proteome sequencing was performed on the same 15 samples used for metabolome and transcriptome analysis. A high Pearson’s correlation coefficient for each group indicated that the data was reliable and could be used for subsequent analyses (Fig. 7a). In total, 27,173 peptides were inferred, and 6642 proteins were confidently identified. The number of peptides in each sample ranged from 17,255 to 23,405, and the number of proteins reached 5319 to 6082 (Supplementary Table 4). A total of 3842 differentially expressed proteins (DEPs) were screened out using the criteria fold change ≥ 1.5, P < 0.05 to indicate upregulated proteins and fold change ≤ 1/1.5, P < 0.05 for downregulated proteins. In the comparisons of S0 vs. S1, S1 vs. S2, S2 vs. S3, and S3 vs. S4, there were 880, 1124, 1870, and 1130 DEPs, including 538, 703, 919, and 300 upregulated DEPs and 342, 421, 951, and 830 downregulated DEPs, respectively (Figure 7b).

**Fig. 7.**
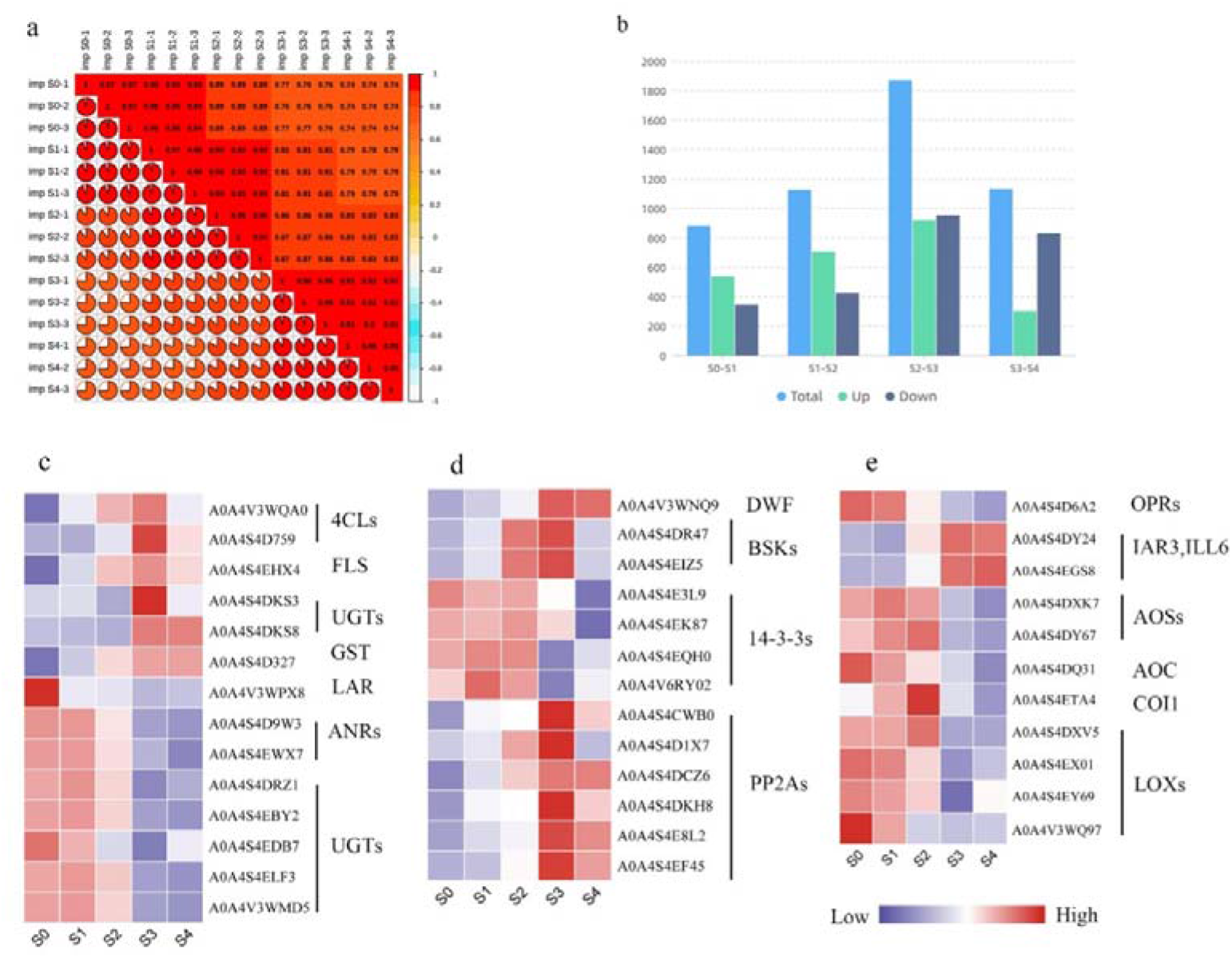
Analysis of proteome data. (a) Correlations of proteome data samples. (b) Histogram of distribution of differential proteins in each period. Heat maps of flavonoid (c), BR-related (d) and JA-related genes (e).

### Differential protein analysis

In the analysis of proteome, many DEPs were found to participate in the flavonoid biosynthesis process, consistent with findings from the transcriptome data, including two 4CLs, one FLS, one GST, one LAR, two ANRs and seven UGTs (Fig 7c). Moreover, many DEPs related to hormones BR and JA (Fig 6d, e) were detected: BR biosynthesis proteins DWF4 (one DEP), BR signal transduction proteins BSK (two DEPs), 14-3-3 (four DEPs), and PP2A (six DEPs); JA biosynthesis proteins LOXs, OPRs, AOSs, AOC; JA receptor protein COI1; JA metabolism-related proteins IAR3/ILL6. Fruthermore, network correlation analysis showed that hormone-related (BR and JA), and flavonoid biosynthesis related proteins and flavonoid metabolites were closely related (Fig. 8), as found in the transcriptome analysis.

**Fig. 8.**
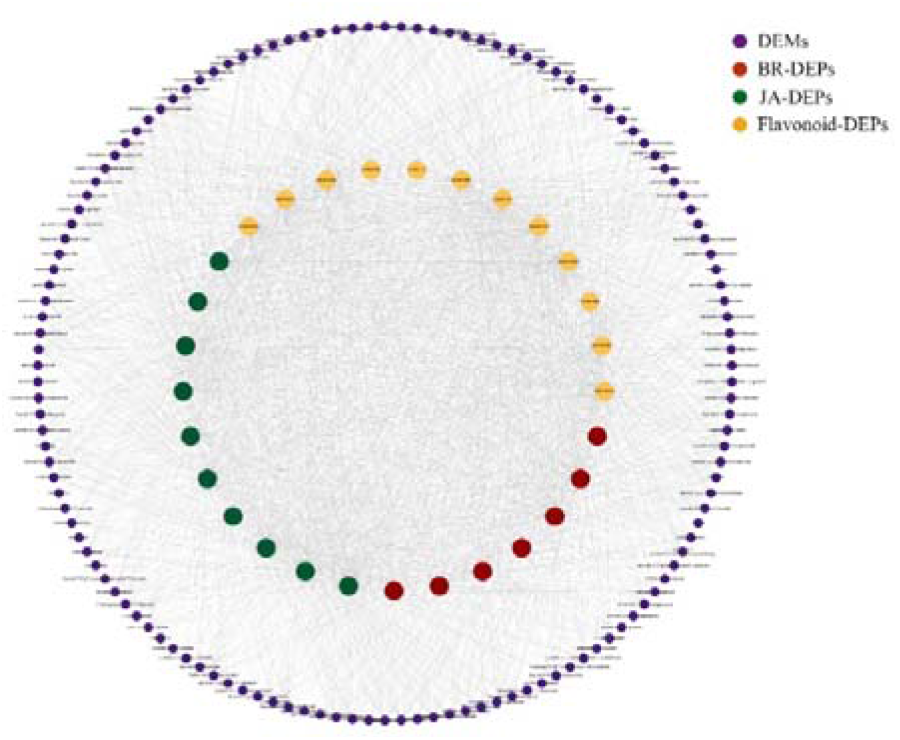
Network correlation analysis of hormone-, flavonoid-related proteins and flavonoid metabolites.

### Conjoint transcriptome, metabolome, and proteome analysis

#### Flavonoid biosynthesis pathway in petals of *C. nitidissima*

Based on the conjoint analysis, DEGs, DEPs and DEMs were mapped to the flavonoid biosynthesis pathway (Fig. 9). The results showed that the expression level of 15 key genes or proteins in flavonoid biosynthesis were significantly different among the five devolpmental stages of petals in *C. nitidissima* and that these differences coincided with the changes in flavonoid content (Fig. 9). The transcript abundance of six key structural genes including *PAL* (five DEGs), *C4H* (four DEGs), *CHI* (three DEGs), *F3H* (one DEGs), *F3’H* (three DEGs) and *F3’5’H* (two DEGs) genes were higher in S3 S4 (golden petal) stages than other stages, consistent with the high abundance of flavonols and petal color change in these stages. For 4CL, four DEGs and two DEPs were detected, and their expression was consistent with the change in flavonol accumulation. For FLS, the key enzyme in the flavonol biosynthesis, there were six DEGs and one DEP identified in the conjoint analysis. UGT (seven DEGs and two DEPs) catalyzes glycosylation of flavonol, and GST (eight DEGs and one DEP), the transpoter of flavonoid, might be responsible for transferring flavonol-glycosides to vacuoles.

**Fig. 9.**
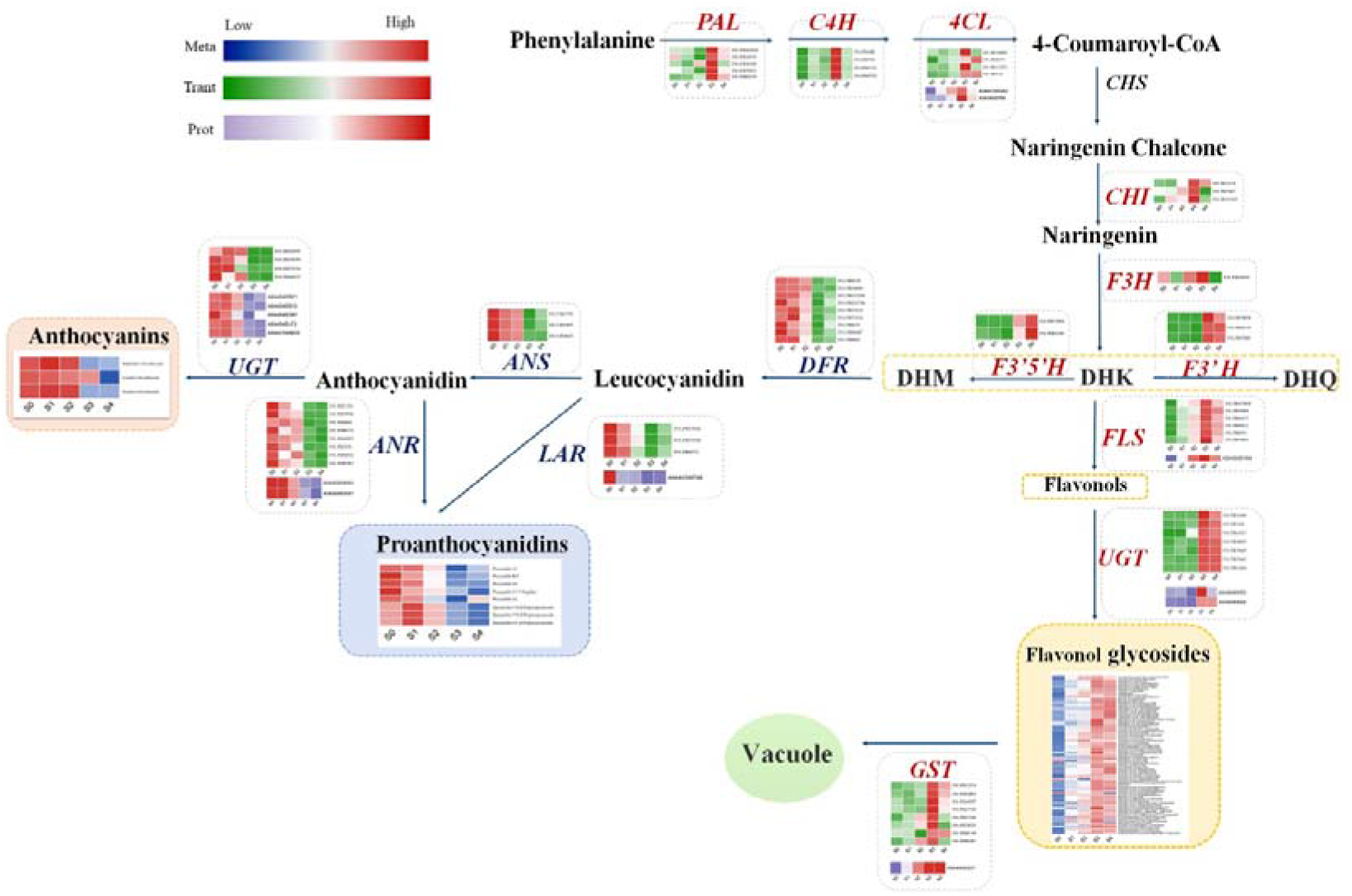
DEGs, DEPs and DEMs involved in the biosynthesis of the flavonoid pathway in *C. nitidissima*.

In the biosynthetic steps from dihydroflavonols (DHK, DHQ and DHM) to anthocyanins and proanthocyanidins, the expression of DFR (nine DEGs), ANS (three DEGs), LAR (three DEGs and one DEP), ANR (eight DEGs and two DEPs), and UGTs (four DEGs and four DEPs) showed a downward trend. This was contrary to the expression of FLS and flavonol acuumulation but consistent with the biosynthesis of anthocyanins and proanthocyanidins.

#### BR and JA may be involved in the regulation of flavonol biosynthesis

The results of the conjoint analysis of DEMs, DEGs, and DEPs suggested that the BR and JA pathways had strong correlations with flavonol biosynthesis (Fig. 6, 8). As shown in Fig. 10, the BR biosynthesis protein DWF4 was upregulated in S3 and S4, consistent with the expression trend of perception and receptor genes *BRI1* (three DEGs) and *BAK1* (12 DEGs). Subsequently, the positive response factors BSK (14 DEGs and two DEPs), BSU1 (three DEGs), and PP2A (24 DEGs and six DEPs) also had increased the expression levels in the gloden petals stages, and the expression of 14-3-3 (six DEGs and four DEPs), a negative regulator of the BR signaling pathway, showed a downward trend in the five petal development stages. Finally, BZR1 (eight DEGs), the core regulator of the BR signaling pathway and responsible for regulating the expression of downstream target genes, showed specifically high expression in the gloden petals stages, consistent with other positive response factors.

**Fig. 10.**
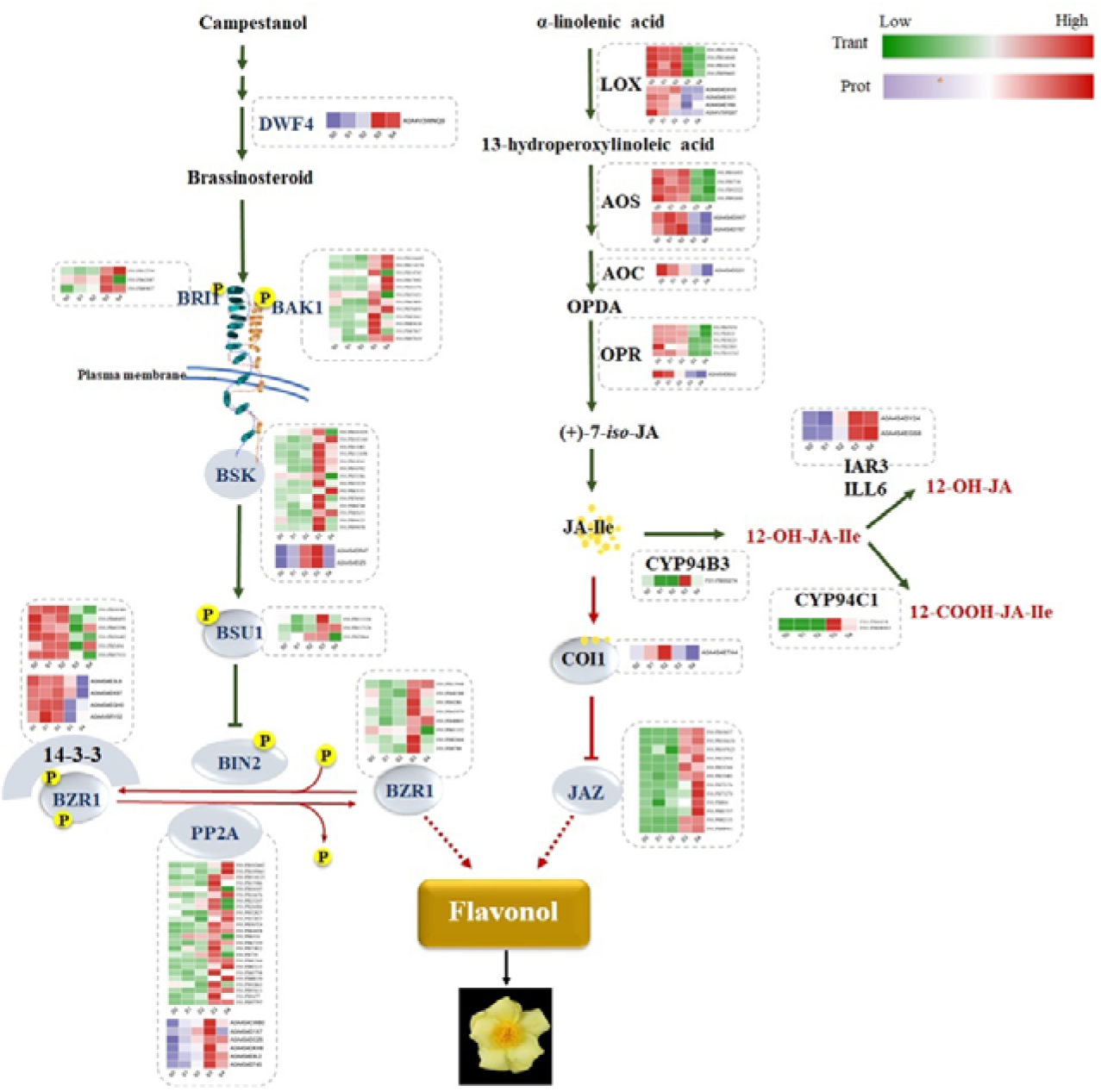
A model for the involvement of BR and JA in the biosynthesis of flavonoids in *C. nitidissima*.

The above results show that the expression of BR-related genes and proteins is positively correlated with flavonol accumulation in *C. nitidissima*. However, the expression of JA-related genes and proteins was negatively correlated with flavonol biosynthesis (Fig. 10). First, the expression level of JA biosynthesis genes OPR, AOC, AOS, LOX was higher in early stages (S0, S1, S2) than in the late stages (S4 and S5). Second, the expression of CYP94B3, CYP94C1, and IAR3/ILL6, which catalyzed JA to form 12-OH-JA or 12-COOH-JA-Ile and then reduced JA signaling, was upregulated in S3 and S4, and the JA receptor protein COI1 was highly expressed in S1 and S2. Finally, the expression of *JAZ* (12 DEGs), the core regulator of JA signaling, showed a upward trend, which was contrary to that of biosynthesis and receptor genes. These results suggest that JA is negatively correlated with flavonol biosynthesis in *C. nitidissima*.

## Discussion

### Flavonol biosynthesis during gloden petal formation in *C. nitidissima*

Flower color is a key ornamental trait of flower plants and an important evalution indicator of the value and quality of flowers. Camellia varities with yellow flowers are scarce (Liu *et al*., 2022), and *C. nitidissima* is a prized woody flower with golden-yellow flowers in camellia, suggesting that it is a valuable resource for molecular mechanism research of yellow flower formation and yellow camellia breeding. In this work, we conducted a flavonoid metabolome analysis of five periods based changes in petal color in *C. nitidissima* (Fig. 1). A total of 323 flavonoids metabolites were obtained, among which flavonol was the most abundant (114 metabolites), accounted for 35.29% (Fig. 2b). Compared with other Camellia plants, the golden petals of *C. nitidissima* show more accumulation of flavonol glycosides, including quercetin-3-O-rutinoside, quercetin-7-O-glucoside, quercetin-3-O-glucoside (Tanikawa *et al*., 2008; Liu *et al*., 2022). Here, we detected 79 flavonol glycosides, including 40 quercetin-related glycosides and 27 kaempferol-related glycosides (Fig. 3), that were positively correlated with golden petal formation. These results suggeted that the glycosides of quercetin and kaempferol are the main flavonoids metabolites in the golden petals of *C. nitidissima*.

The biosynthetic pathway of plant flavonoids has been elucidated (Winkel-Shirley, 2001; Dong & Lin, 2020; Liu *et al*., 2021). The expression of of flavonoid structural genes, such as *PAL*, *4CL*, *CHI*, *CHS, FLS, F3’H, DFR, ANS, LAR*, and *ANR* produces various flavonoid metabolites, including flavonols, anthocyanins and proanthocyanidins (Lepiniec *et al*., 2006; Sasaki & Nakayama, 2015; Xu *et al*., 2020). In this work, transcriptome and proteome sequencing were implemented to explore the regulatory pathways of flavonol acuumulation. First, the flavonol biosynthesis genes *PAL*, *4CL*, *C4H, CHI, F3H, F3’5’H, F3’H, UGT, GST*, and the key limiting gene *FLS*, were found to be highly expressed in the gloden petal phase (Fig 9), consistent with the accumulation of flavonol glycosides. Second, the expression of anthocyanin and proanthocyanidin biosynthesis genes *DFR, ANS, UGT, LAR, ANR* showed a downward trend during petal development (Fig 9). *FLS* and *DFR* are key structural genes by which flavonoids enter different synthetic branches (Kubra *et al*., 2021; Liu *et al*., 2021; Wang *et al*., 2022), and overexpression of *FLS* increases the biosynthesis of flavonols (Park *et al*., 2020; Yu *et al*., 2021). Heterologous expression of *Rosa rugosa RrDFR1* and *Petunia hybrida PhDFR* in tobacco has been reported to inhibit the expression of the endogenous *NtFLS* and promote the accumulation of anthocyanins (Luo *et al*., 2016). Studies have also shown that DFR and FLS competed for common dihydroflavonol substrates, and their exprssion inhibits each other’s transcription (Jiang *et al*., 2020; Chen *et al*., 2021). In this work, during the early development stage (S0-S2) of *C. nitidissima, DRF* was highly expressed, and flavonoids entered the anthocyanins and proanthocyanidins branching pathway. When petals entered the golden stage (S3-S4), expression of *DFR* was downregulated, and *FLS* was highly expressed, and flavonols were synthesized. In addition, the expression of proteins 4CL, FLS, GST, LAR, ANR and UGT was consistent with their gene’s expression and with flavonols biosynthesis in *C. nitidissima*. Thus, the upregulated expression of flavonol-related genes and proteins, especially FLS, and the downregulated expression of anthocyanin and proanthocyanidin genes lead to massive biosynthesis of flavonol glycosides in the process of golden color formation.

bHLH and MYB transcription factors are the vital transcription regulator of flavonoid metabolism (Li *et al*., 2019; Sun *et al*., 2020a; Rajput *et al*., 2022). GtMYBP3 and GtMYBP4 regulate early biosynthesis genes of flavonoids, *FNS* and *F3’H*, and promete flavonol biosynthesis (Nakatsuka *et al*., 2012). AcB2, a bHLH transcription factor, had been reported to interact with AcMYB1 and increase anthocyanin biosynthesis in onion (Li *et al*., 2022). After our KEGG analysis, there were 20 *bHLH* and six *MYB* genes annotated into the flavonoid pathway (Fig 5b, c). Furthermore, network correlation analysis showed that 16 *bHLH* and five *MYB* genes were closely related to flavonoids, hormone genes, and flavonoid metabolites (Fig. 8), indicating that these *bHLH* and *MYB* genes might be key regulators of flavonoid pathway in *C. nitidissima*.

### BR and JA may be inloved in the regulation of flavonol biosynthesis during golden petal formation

Hormones are crucial for the processes of plant growth and development, including flavonoid metabolism (Dong & Lin, 2020; LaFountain & Yuan, 2021). Such hormones include auxin (Wang, YC *et al*., 2018), JA (Ni *et al*., 2020), GA (Tan *et al*., 2019), BR (Liang *et al*., 2020), strigolactone (Wang *et al*., 2020), ethylene (Ma *et al*., 2021) and abscisic acid (Loreti *et al*., 2008). In this study, many BR and JA biosynthesis and signaling genes or proteins were detected (Fig. 5). In *A. thaliana*, BR signaling regulator BES1 (also named BZR2) was shown to directly bind to the promoters of *AtMYB11, AtMYB12*, and *AtMYB111* and represses their expression, thereby reducing flavonol biosynthesis (Liang *et al*., 2020), which indicated that BR negatively regulated flavonol biosynthesis. However, the expression of *BZR1* was positively correlated with flavonol accumulation in *C. nitidissima*, the same as for other BR biosynthesis and signaling genes or proteins (Fig. 10). Network correlation analysis showed that genes and proteins of the BR pathway were closely related to the MYB and bHLH transcription factors, flavonoid pathway, and flavonoid metabolites (Fig. 6, 8). These results suggest that BR positively regulates flavonol biosynthesis in *C. nitidissima*, the opposite situation to that found in *A. thaliana*.

JA positively regulates biosynthesis of various flavonoids, including anthocyanins (Premathilake *et al*., 2020), proanthocyanidins (An *et al*., 2021; Zhu *et al*., 2022), flavonols, and flavones (He *et al*., 2022). MeJA was shown to promote the flavonol biosynthetic enzyme genes *FLS*, *F3H, CHS*, and *CHI* and transcription factor *MYB81* in *Gynostemma pentaphyllum* (Huang *et al*., 2022), and MeJA treatment increased the flavonol content in red raspberry (Flores & Ruiz del Castillo, 2014). JAZ, a repressor of the JA signaling, interacted with MYB or bHLH (members of MBW complex) and inhibited the flavonoids biosynthesis (Yan *et al*., 2021). And CtJAZ1 interacted with CtbHLH3 to inhibit MBW formation and consequently inhibit flavonol and flavone biosynthesis in safflower (He *et al*., 2022). Thus, previous studies indicat that the JA pathway is positively related to flavonol biosynthesis in plants. However, in this study, the expression of JA biosynthesis, metabolism and signaling genes or proteins was negatively related to the level of flavonol accumulation (Fig. 10). In *C. nitidissima*, the BR and JA hormone pathways might be involved in the flavonol metabolism, with the functions of BR and JA being different from those in other plants; this requires further study.

BZR1/2 and JAZ, the core regulators of BR and JA, are the ‘bridges’ by which BR and JA participate in the regulation of plant growth and development. BR regulates flavonol biosynthesis via BES1 (BZR2)-MYBs pathway (Liang *et al*., 2020), and CtJAZ1 interacts with CtbHLH3 to repress flavonol biosynthesis (He *et al*., 2022). Thus, BR and JA might regulate flavonol metabolism through interactions between BZR1/2 and JAZ and MBW complex members (MYBs and bHLHs). In this study, the expression trends of *BZR1* and *JAZ* were consistent with those of multiple *MYB* and *bHLH* genes (Fig. 5). Furthermore, network correlation analysis showed that *BZR1* and *JAZ* genes were closely related to the *MYB* and *bHLH* genes (Fig. 6). These results suggest that BR and JA might be involved in the regulation of flavonol biosynthesis via the BZR1 or JAZ-MYB or bHLH pathway; however, this requires further study.

## Supplementary data

Supplementary Table1. The primers sequence.

Supplementary Table 2 Summary of reads after sequencing.

Supplementary Table 3. The results of functional annotation of unigenes.

Supplementary Table 4 Summary of sequencing proteins.

Supplementary Fig. 1. The analysis of differential expressed metabolites (DEMs).

Supplementary Fig. 2. Annotated drawing with NR database.

Supplementary Fig. 3 RT-PCR analysis of the 14 candidate genes.

Supplementary Fig. 4 Heat map of bHLH (a) and MYB (b) genes.

## Author contributions

Y.F.: Investigation, Formal Analysis, Writing—original draft. J.L.: Funding acquisition. X.C.: Investigation. H.Y.: Writing—review and editing. Z.F.: Visualization. S.Y.: Investigation. M.W.: Investigation. X.L.: Validation. W.L.: Conceptualization, Formal Analysis, Writing—review and editing. All the authors reviewed and approved the final manuscript. All authors have read and agreed to the published version of the manuscript.

